# Feeding Drosophila highly radioresistant fungi improves survival and gut morphology following acute gamma radiation exposure

**DOI:** 10.1101/2025.07.10.664168

**Authors:** Robert P. Volpe, Hye Jin Hwang, Rachel T. Cox

**Affiliations:** Department of Biochemistry and Molecular Biology, Uniformed Services University, Bethesda, MD, 20814; Henry M. Jackson Foundation for the Advancement of Military Medicine, Bethesda, MD, 20817

**Keywords:** fungi, radioresistant, irradiation, lifespan, gut, *Drosophila*

## Abstract

Diverse fungi have been historically vital reservoirs of drug discovery, providing life-saving pharmaceuticals. Many species of fungi, yeasts in particular, are highly resistant to radiation, with their cellular contents potentially conferring dietary radioresistance. We developed a *Drosophila* model to test whether feeding two highly radioresistant fungi, *Aureobasidium pullulans* and *Rhodotorula taiwanensis,* could improve fly lifespan and gut morphology after acute irradiation. We constructed a dosimetry curve for the lifespan response of males and females to irradiation and found dose-dependent and sex-specific effects on lifespan. We also determined that the sex-specific response to irradiation correlated with nuclear morphology defects in the gut, with the more radiosensitive males displaying increased midgut cellular holes and aberrant nuclear morphology. To determine if feeding *Aureobasidium pullulans* and *Rhodotorula taiwanensis* before irradiation could improve survival and gut morphology, we first exclusively fed males and females each fungus and observed that they tolerated the diet well. Using these methods, we found that only two days of prefeeding *Aureobasidium pullulans* increased male lifespan, but not female, after irradiation, and improved nuclear morphology in the gut. However, dietary *Rhodotorula taiwanensis* was not protective. Overall, this study identified a highly radioresistant dietary fungus, *Aureobasidium pullulans*, as effective for extending male *Drosophila* lifespan and improving gut morphology following irradiation. Since the gut is particularly sensitive to the effects of irradiation, this fungus indicates a potential therapeutic for patients undergoing radiotherapy. Furthermore, this method could identify additional radioresistant fungi that protect the gut from radiation injury.

## Introduction

Fungi synthesize a large number of naturally occurring bioactive molecules from which researchers have identified a wide array of vital pharmaceuticals such as antibiotics, immunosuppressants, and cholesterol-lowering drugs (Aly et al., 2011; Prescott et al., 2023). Within this large and diverse kingdom are species that are highly radiation-resistant, with fungal species growing in the reactor of the damaged Chernobyl Nuclear Power Plant among the first ones identified (Zhdanova et al., 1994). Since then, many species of fungi have been shown to survive acute and chronic exposures to astoundingly high levels of ionizing radiation, far beyond any that are naturally occurring (Shuryak et al., 2019; Tkavc et al., 2017). Several mechanisms contribute to the astonishing ability of these microbes to grow and reproduce in highly irradiated environments. Some radiation-resistant organisms accumulate antioxidant compounds such as manganous peptide phosphate complexes to survive the oxidative stress of IR (Gaidamakova et al., 2022; Sharma et al., 2017). Melanin often accumulates in response to stress in various fungi, and its chemical properties include broad optical absorption and the ability to interact with ionizing radiation, lending it antioxidant activity (Cordero et al., 2017; Dadachova & Casadevall, 2008; Zhdanova et al., 1994). Given the diversity of the kingdom and the fact that so many species demonstrate radioresistance to levels of IR not found in nature, this suggests that additional metabolites and molecules conferring radioresistance have yet to be identified.

Using *Drosophila*, we investigated whether ingestion of two radioresistant fungi, *Aureobasidium pullulans* (*A. pullulans*) and *Rhodotorula taiwanensis* (*R. taiwanensis*), could provide radioprotection. *A. pullulans*, a dimorphic ascomycetous yeast, grows on the inner parts of the damaged Chernobyl Nuclear Power Plant and the International Space Station, and is considered a polyextremotolerant fungus (Campana et al., 2022; Checinska Sielaff et al., 2019; Zhdanova et al., 1994). *A. pullulans* is usually light in color but becomes darker under stress due to melanin production, which is thought to confer enhanced survival (Campana et al., 2022; Dadachova et al., 2007). *R. taiwanensis*, a basidiomycetous carotenogenic yeast, was characterized for its ability to tolerate high levels of chronic radiation and low pH as part of an initiative to identify fungi that could be harnessed for the bioremediation of radioactive waste sites (Tkavc et al., 2017). *R. taiwanensis* can grow under chronic 66 Gy/h gamma radiation at pH 2.3, giving this fungus remarkable resiliency (Tkavc et al., 2017). Since *R. taiwanensis* and *A. pullulans* survive IR in the absence of melanin, there are likely unidentified metabolites and small molecules that function to protect them against the damaging effects of IR.

We hypothesized that ingestion of *A. pullulans* or *R. taiwanensis* could provide radioprotection to Drosophila. *Drosophila* have a rapid lifecycle, short lifespan, and large genetic toolkit, making them an ideal animal in which to test for dietary protection against IR. Furthermore, *Drosophila* have organ systems, including the gut, that are highly homologous to human physiology, sufficient for modeling numerous human diseases (Bier, 2005; Verheyen, 2022). Perhaps most important to our inquiry, *Drosophila* are natural fungivores and readily consume diverse yeast species with notable effects on physiology and behavior (Anagnostou et al., 2010; Cooper, 1960). Thus, natural feeding would allow direct contact between the radioresistant fungi and highly radiosensitive gut cells, optimizing any potential benefits.

To test the protective effects of dietary *A. pullulans* and *R. taiwanensis* after acute exposure to gamma radiation, we first constructed radiation survival curves for male and female lifespans to identify doses that were not acutely detrimental to fly health, yet high enough to make lifespan measurements manageable. Males were more sensitive to the effects of IR compared to females, as expected. In addition, we observed that male gut morphology was significantly more disrupted compared to females receiving same IR dose. Next, we fed males and females exclusive diets of both fungi and determined that, although they shortened lifespan, neither was acutely toxic, nor did they have detrimental effects on development. To test for dietary protection, we fed flies either fungus for two days before IR exposure and determined that *A. pullulans*, but not *R. taiwanensis*, enhanced radiation survival of males, but not females, for the next fifteen to twenty days. Males fed *A. pullulans* and subsequently exposed to IR showed improved nuclear morphology in the gut compared to non-fed control males, suggesting this could be a contributing factor to the increased survival. Overall, this study demonstrates that *Drosophila* is an effective model to successfully identify highly radioresistant dietary fungi that confer radioresistance to the host. Notably, we identified *A. pullulans* as a promising radioprotectant, which could be further tested in higher eukaryotes.

## Materials and methods

### Drosophila and fungal strains

Fly strains used in this study: *w^1118^* and *catalase^n1^*/*TM3, Sb^1^, Ser* (Kyoto *Drosophila* Stock Center, #107554). Fly stocks were maintained in standard cornmeal fly food vials at room temperature and a 12-hour light cycle. The strain of *Aureobasidium pullulans var. namibiae* (*A. pullulans*) (EXF-1147) used in this study was obtained from Microbial Culture Collection EX curated by the Biotechnical Faculty of the University of Ljubljana in Ljubljana, Slovenia. The strain of *Rhodotorula taiwanensis* (*R. taiwanensis*) (MD-1149) was isolated from an acid mine drainage facility in Allegany County, Maryland, USA, and curated in the collection of Dr. Michael Daly of the Department of Pathology of the Uniformed Services University of the Health Sciences in Bethesda, Maryland, USA. Dietary yeast cultures were grown at room temperature on standard YPD agar plates [1% Bacto Yeast Extract (Cat# 212750, Thermo Fisher Scientific, Waltham, MA, USA), 2% Peptone (Cat# 211677, Thermo Fisher Scientific, Waltham, MA, USA), 2% Dextrose (Cat# PHR1000, Millipore Sigma, St. Louis, MO, USA), 2% Bacto Agar (Cat# DF-0140-01-0, Thermo Fisher Scientific, Waltham, MA, USA)] covered with gamma-sterilized dialysis membrane (Frey Scientific Dialysis Tubing, Cat# 1591654, Dialysis Tubing, Frey Scientific®, Greenville, WI, USA) to ensure only yeast was removed during collection. For pre-feeding IR exposures of cultures, *A. pullulans* and *R. taiwanensis* were grown for 4 days and 1 day, respectively, under chronic irradiation at 35 Gy/hr in a ^137^Cs-sourced gamma irradiator (Shepard and Associates 68A Mark1, San Fernando, CA, USA). Unirradiated fungi cultures were grown in the same location outside the irradiator to ensure both cultures grew under otherwise identical conditions. Acute radiation tolerances (D_10_) for both strains were determined by incremental irradiation of liquid YPD culture (OD_600_ = 0.8) in a ^60^Co-sourced gamma irradiator (Shepard and Associates, San Fernando, CA, USA) followed by colony forming unit (CFU) assay on solid YPD medium as previously described (Daly et al., 2004). To effect a bisecting exposure of *A. pullulans* culture, we cast a cylindrical radiation shield of pure lead (Rotometals, Inc., San Leandro, CA, USA) with a central cavity of approximately 65 mm x 30 mm x 12 mm to snugly fit a standard 60 mm petri dish. *A. pullulans* was streaked on solid YPD agar plate, placed in the radiation shield leaving half the plate exposed, and irradiated at ∼35 Gy/hr for 24 hours in a ^137^Cs-sourced irradiator. The partially irradiated culture was then allowed to recover for 3 days. To observe phenotypic switching of melanin expression in response to cyclical irradiation, 20 µL of *A. pullulans* in liquid YPD (OD_600_ 0.8) was aliquoted to the center of a YPD agar plate. After one day of growth, the culture was exposed to alternating cycles of chronic radiation at 35 Gy/hr for 2 days followed by 2 days without exposure. To visualize accumulation of melanin in the cell walls of *A. pullulans* hyphae, cultures were imaged using a Motic BA210E (Motic, Universal City, TX, USA) at 400x (40x objective x 10x eyepiece).

### Fungal feeding

For administration of fungal diets, flies were housed in feeding chambers made from agar feeding plates encapsulated by perforated plastic beakers. Aliquots of fungi were carefully harvested with a silicone spatula from 2-3 day-old cultures grown on YPD plates and added to agar plates. Flies were allowed to feed for two days, irradiated, then transferred to standard food vials supplemented with a small amount of live dry yeast (*Saccharomyces cerevisiae* (*S. cerevisiae*) (Red Star Active Dry Yeast, Red Star Yeast Company LLC, Milwaukee, WI, United States)). For determining the toxicity of, and survivorship on, *A. pullulans* and *R. taiwanensis*, diets, twenty males or females, in triplicate, were continuously fed fungal paste for the entire lifespan, changing the plate and fungal paste every 1-2 days. A paste of water and *S. cerevisiae* was used as a control. To determine pupation and eclosion rates, twenty first instar larvae collected one day after egg laying were transferred into agar plates supplemented with *A. pullulans*, *R. taiwanensis* or a paste of control *S. cerevisiae* and grown at room temperature.

The number of pupae was counted every 24 hours after the onset of pupation. The number of eclosed adults was counted each day after the onset of eclosion. Each experiment was performed in triplicate. To visualize consumption of the fungi, representative flies and larvae were imaged using an Accu-scope 3076 digital microscope 0.67x-4.5x (Accu-Scope, Commack, NY, USA).

### Irradiation, dosimetry, and lifespan analysis

Flies were exposed to gamma radiation in standard polystyrene *Drosophila* vials (Genesee Scientific, Cat# 32-109, Morrisville, NC, USA) capped with cellulose-acetate stoppers (Genesee Scientific, Cat# 49-102, Morrisville, NC, USA) containing standard cornmeal fly media (8.25% corn syrup (Karo Light Corn Syrup, ACH Food Companies, Oakbrook Terrace, IL, United States), 5% corn meal (Cat# 62-100, Genesee Scientific, El Cajon, CA, United States), 1.25% dry inactivated yeast (Cat# 62-103, Genesee Scientific, El Cajon, CA, United States), 0.75% soy flour (Cat# 62-115, Genesee Scientific, El Cajon, CA, United States), 0.5% propionic acid (Cat# P1386, Millipore Sigma, St. Louis, MO, United States), 0.1% Tegosept (Ca# 20-258, Genesee Scientific, El Cajon, CA, United States) with 0.04 g live dry yeast (Red Star Active Dry Yeast, Red Star Yeast Company LLC, Milwaukee, WI, United States). Six vials, two sets of matching triplicates, were irradiated per radiation treatment using a vendor-calibrated, ^137^Cs-sourced gamma irradiator. Vials were placed on a rotating platform to ensure uniform exposure of all samples to the source emission. Consistent and precise dosimetry for all sample irradiations was achieved by calculating the length of each exposure, accounting for daily source decay using the standard decay equation. For lifespan analysis, twenty male and female adults were monitored daily after irradiation for their full lifespan, which was performed minimally in triplicate as described in *Fungal feeding* above. Since irradiation significantly weakens the animals, the determination of death for each fly was made by observing whether it failed to respond repeatedly to gentle probing, as indicated by head twitching and/or leg movements. To perform statistical analysis, the average daily survivorship data from the triplicates were compiled and entered into the Online Application for Survival Analysis 2 (OASIS) (https://sbi.postech.ac.kr/oasis2/) (Han et al., 2016). Statistical significance between control and experimental conditions was determined in OASIS using the Wilcoxon-Breslow-Gehan test.

### Immunostaining and gut injury analysis

Post-irradiated day 2 adult flies were dissected and immunostained as previously described, with slight modifications (Chen & Johnston, 2022). Briefly, day 2 adult flies were irradiated and subsequently maintained in standard food vials for an additional two days. The guts were dissected in 1x phosphate-buffered saline (PBS) and fixed with 4% formaldehyde in 1x PBS for thirty minutes at room temperature. After washing with Antibody wash solution (AWS, 0.1% TritonX-100, and 1% BSA in PBS) three times for twenty minutes, the tissues were stained with Alexa Fluor^TM^ 488 Phalloidin (Cat# A12379, Invitrogen, Waltham, MA, USA) overnight at 4 ^°^C. Following washes with AWS twice for 20 minutes, tissues were stained with 4ʹ,6-Diamidino-2- phenylindole (DAPI) for 10 minutes at room temperature and mounted in Vectashield Antifade Mounting Medium (Cat# H-1000, Vector Laboratories, Newark, CA, USA). Images were obtained using a Zeiss LSM 980 confocal laser scanning microscope with a 63x objective lens (Carl Zeiss Microscopy LLC, White Plains, NY, USA). The R4 region of the midgut was chosen, and a z-stacked image was obtained for the whole layer of enterocytes. The total number of guts analyzed is indicated in each figure legend. Quantification of abnormal nuclear shape, disrupted cellular barriers, and holes in the actin filament layer of enterocytes was determined by subjective measurement. Abnormal nuclear shape was determined if a nucleus had a smaller size and a loss of staining in the nucleus. Disrupted cellular barriers between enterocytes were evaluated by loss of actin and abnormal cellular shapes, as demonstrated by the actin staining. Holes in an actin filament layer were counted if the following conditions were satisfied: at least five clear round-shaped holes with a diameter of more than 1.5 µm in an actin layer within a single cell, or two holes with a diameter of more than 2.5 µm. The percentage of animals with gut injury in each replicate was calculated, averaged, and plotted in a graph generated using GraphPad Prism (GraphPad Software, version 10.4.2, Boston, Massachusetts USA, www.graphpad.com). Significant differences between groups were tested as described in *Statistical analyses* below.

### Statistical analyses

To quantify the relationship between radiation dose and mortality, linear regression was conducted in Microsoft Excel (version 16.94.1), and the coefficient of determination (R²) was reported as a measure of goodness-of-fit. Lifespan data were analyzed using the OASIS 2 online platform for survival analysis (Yang et al., 2011; https://sbi.postech.ac.kr/oasis2/) using compiled average of three triplicates. Mean lifespan statistical significance between groups was determined using log-rank and weighted log-rank (Wilcoxon–Breslow-Gehan) test (Landes et al., 2012). The analysis of gut injury was performed using GraphPad Prism (GraphPad Software, version 10.4.2, Boston, Massachusetts USA, www.graphpad.com). Two-way analysis of variance (ANOVA) with multiple comparisons followed by Tukey’s post hoc test was performed to evaluate the effect of sex (male, female) and irradiation (-IR, +IR) on abnormal physiological events. Mean lifespans, standard errors, and Bonferroni p-values were reported to assess the impact of dietary intervention on survival following irradiation and are presented in Supplementary Table 1.

### Biological, chemical, and radiological safety

All experimental work was performed in strict adherence to the biological, chemical, and radiological safety protocols and regulations established by the Uniformed Services University of the Health Sciences and its Environmental Health and Radiation Safety Divisions.

## Data availability statement

All data are contained within the manuscript.

## Results

### *Drosophila* exhibit dose-dependent lifespan sensitivity to exposure to acute irradiation

In order to use *Drosophila* as a model to test dietary radioprophylactic properties of highly radioresistant fungi, we designed a protocol for irradiating adult flies and determined lifespan dose curves for a range of radiation exposures. For consistent irradiation, six standard fly food vials were elevated to the appropriate height on a rotating platform in a ^137^Cs-source irradiator (Fig. 1A). A blank vial was placed in the middle of the vial cluster. This configuration ensured that all flies were exposed to the correct dose of irradiation. This also enabled us to simultaneously test experimental and control groups in triplicate. The normal (*w^1118^*) *Drosophila* lifespan is approximately 100 days (Fig. 1B, black lines). We tested the effect of 0-1500 Gy gamma radiation on the lifespan of *w^1118^* male and female adult flies (Fig. 1B). Twenty newly eclosed males and females, in triplicate, were simultaneously exposed to increasing doses of acute gamma radiation (Fig. 1B). Exposed males (Fig. 1B, dashed lines) were more sensitive to the effect of IR compared to females (Fig. 1B, solid lines) for every dose. In addition, the exponential decline in the number of days to reach LD_50_ of males and females was observed with increased IR dose, with a strong coefficient of determination (R^2^) values of 0.9728 and 0.9095 for males and females, respectively (Fig. 1C). Finally, we tested whether *Drosophila* mutants lacking the free-radical scavenging enzyme *catalase* (*cat*) are more sensitive to IR exposure using our method. Day 2 adult *cat^n1^* null mutants exposed to 1000 Gy had shorter lifespans compared to *w^1118^*, with males more sensitive than females (Fig. 1D, p = 0.0002 (***) (female) and p=0.000049 (****) (male)). Together, these data demonstrate that our method exposing male and female *Drosophila* to IR consistently shortened lifespan in a dose-sensitive and sex-specific manner.

**Fig. 1:**
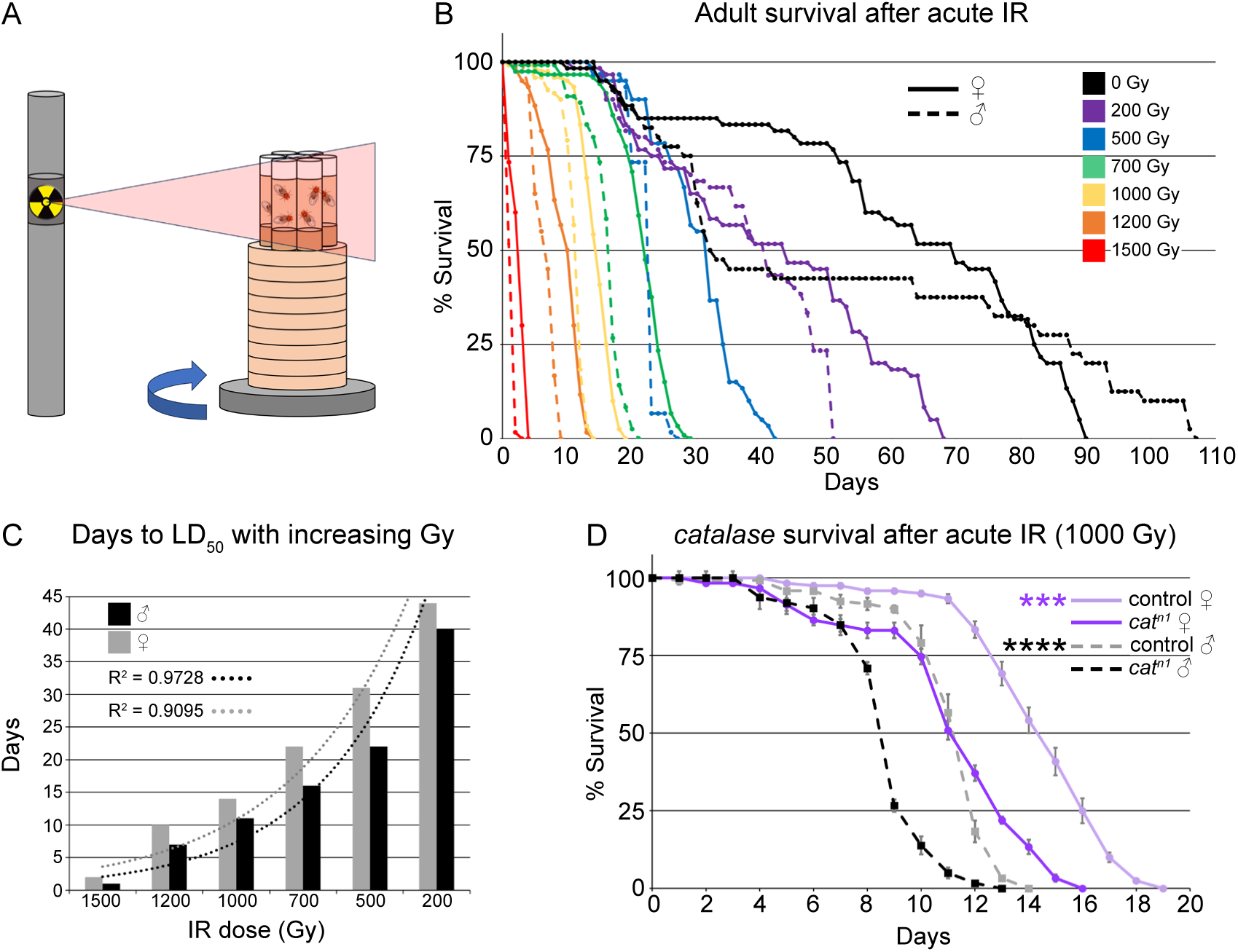
Dosimetry and radiation sensitivity of *Drosophila*. (A) Schematic of *Drosophila* irradiation sequence. (B) Lifespan of male and female flies acutely exposed to 0-1500 Gy gamma radiation. Radiation survival negatively correlates with dose. Males (dashed lines) were more sensitive than females (solid line) for all doses. (C) Graph showing a negative exponential relationship between the days to LD_50_ for males and females and increasing IR dose, indicated by R^2^ close to 1.0 for both sexes. (D) Lifespan of control (*w^1118^*) and *catalase^n1^* (*cat^n1^*) mutants after 1000 Gy acute irradiation. *cat^n1^* null flies are more sensitive to irradiation than control, and males are more sensitive than females for both genotypes. IR = ionizing radiation. Graphs were plotted using Microsoft Excel. Each point on the lifespan represents the average of triplicates of twenty flies that were irradiated simultaneously. Significance: (D) p = 0.0002 (***) for females and 0.000049 (****) for males.

### *Drosophila* male guts are more sensitive to the effects of irradiation compared to females

As a single-layered epithelium, the adult fly midgut responds quickly to environmental changes, including nutrient fluctuations and tissue damage, in order to maintain homeostasis. Given this rapid physiological response, we conducted immunostaining on dissected midguts from flies exposed to 1000 Gy to identify physiological differences between males and females (Fig. 2, Fig. S1). To do this, we labeled the midgut enterocytes to visualize actin filaments and nuclei, then imaged the R4 region of the midguts using confocal microscopy. Irradiated males had abnormal nuclear shape (Fig. 2B,B”, E, Fig. S1D,D”,E,E”,F,F”, arrowheads), disrupted cellular barriers (Fig. 2B,B’,F, Fig. S1D,D’,E,E’), or visible holes within actin alignments (Fig. 2G, Fig. S1D,D’,E,E’, asterisks) compared to non-irradiated males that had normal nuclear shape (Fig. 2A,A”,E, Fig. S1A,A”,B,B”,C,C”) and well-organized actin filament distribution (Fig. 2A,A’,F, Fig. S1A,A’,B,B’,C,C’). In contrast, irradiated females had normal nuclear shape (Fig. 2D,D”,E, Fig. S1J,J”,K,K”,L,L”) and fewer holes within the actin cytoskeleton (Fig. 2D,D’,G) compared to irradiated males. However, irradiated females had significant numbers of enterocytes with indistinct cellular barriers compared to non-irradiated females (Fig. 2D,D’,F, Fig. S1J,J’,K,K’,L,L’). This suggests that the enterocytes of male guts are more vulnerable to the effects of IR compared to female guts, which could contribute to the difference between male and female survival after irradiation.

**Fig. 2:**
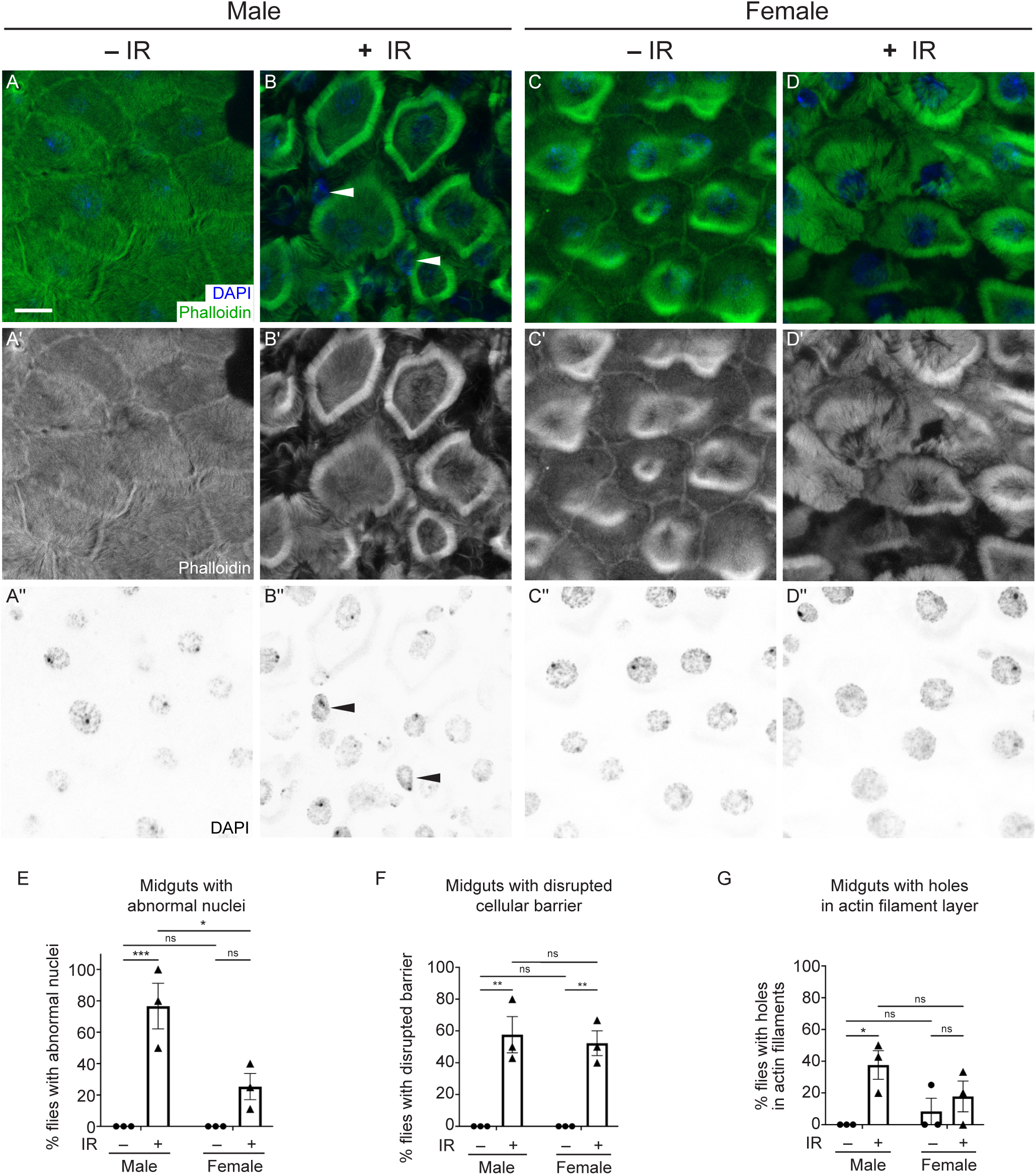
*Drosophila* male guts are more sensitive to the effects of irradiation compared to females. (A-D”) Dissected midguts from Day 4 adults exposed to IR (+IR) at day 2 and unexposed controls (-IR) labeled with phalloidin to label actin filaments and DAPI to label nuclei. The images of the R4 regions of midguts from each experimental condition were obtained using a confocal microscope with z-stacked for a layer of enterocytes. (A-A”) The enterocytes of a representative non-irradiated male control showing clear cellular barriers (A,A’) and good shape of nuclei (A,A”). (B-B”) The enterocytes of a representative male two days after irradiation showing abnormal nuclear shape (B, B”, arrowheads) and ambiguous cellular morphology due to disrupted cellular barriers (B,B’). (C-C”) The enterocytes of a representative female control and (D-D”) a representative female two days after irradiation showing normal nuclear morphology (C,C”,D,D”) compared to irradiated males (E). Irradiated females had disrupted cellular barriers between enterocytes (D,D’,F) similar to irradiated males (B,B’,F). (E-G) Each graph indicates the percentage of animals with guts showing abnormal nuclei (E), (disrupted cellular barriers (F), and holes in the actin filament layer in enterocytes (G). The quantification method is described in the Materials and Methods section. Each point on the graph represents the percentage of each replicate, and each bar represents the arithmetic mean of triplicates with a standard error of the mean (SEM). Data were analyzed statistically using a two-way ANOVA with multiple comparisons, followed by Tukey’s post hoc analysis. The following numbers of the midguts were assayed per experiment: male (-IR) (6, 7, 7); male (+IR) (5, 7, 4); female (-IR) (4, 8, 9); female (+IR) (5, 8, 9). Significance of multiple comparisons: (E) p = 0.0009 (***) for males and 0.0108 (*) for male (+IR) vs female (+IR); (F) p = 0.0016 (**) for males and 0.0030 (**) for females; (G) p = 0.0379 (*) for males. ns = not significant. (A,B,C,D) Green = phalloidin, blue = DAPI. (A’,B’,C’,D’) White = phalloidin. (A”,B”,C”,D”) White = DAPI. (A-G) -IR: non-irradiated, +IR: 1000 Gy irradiation. Scale bar: 10 μm in A for A-D”.

### Rhodotorula taiwanensis and Aureobasidium pullulans are highly radioresistant fungi

*R. taiwanensis* and *A. pullulans* (var. *namibiae*) are highly radioresistant fungi and potential candidates for dietary prevention strategy to injury from acute IR (Fig. 3A) (Liu et al., 2017; Tkavc et al., 2017). *R. taiwanensis*, a basidiomycetous carotenogenic fungus, is radioresistant to acute exposures of 2,500 Gy and chronic exposures of 60 Gy/hr (Fig. 3A,B, (Tkavc et al., 2017)).

**Fig. 3:**
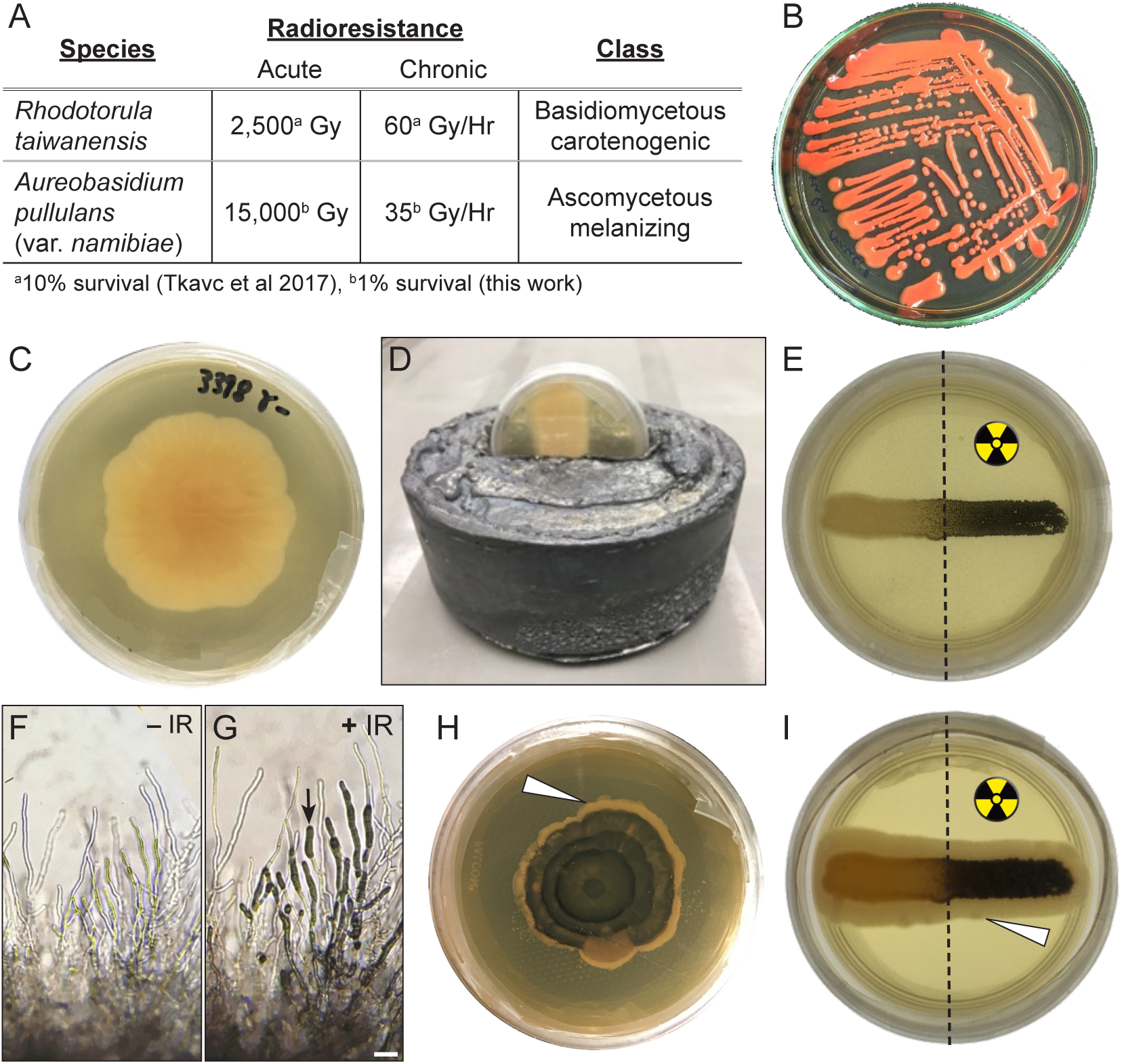
*Rhodotorula taiwanensis* and *Aureobasidium pullulans* are highly radioresistant fungi. (A) Characteristics of *R. taiwanensis* and *A. pullulans*. Both are highly radioresistant. (B) Agar plate showing streaked red *R. taiwanensis.* (C) Agar plate with a colony of white *A. pullulans.* (D) Lead shield used for irradiation experiments on an agar plate. Only the top half of the plate is exposed to irradiation. (E) Culture streak of *A. pullulans* protected (left) and unprotected (right) from exposure to irradiation. The right half of the culture is black due to melanin production following irradiation. (F, G) Micrographs of *A. pullulans* hyphae without (F, −IR) and with (G, +IR) irradiation. Melanin deposits (arrow) are visible in the hyphae. (H, I) Agar plate (H) and colony streak (I) of radiation-induced melanized *A. pullulans* allowed to grow without radiation. New fungal growth after irradiation is non-melanized and white (arrow heads). The colony shown in I is the same as the colony in E. Scale bar: 50 μm in G for F & G.

*A. pullulans*, a dimorphic ascomycetous fungus, normally grows as a white colony (Fig. 3C). We assessed that *A. pullulans* is radioresistant to acute exposures as high as 15,000 Gy and chronic exposures of 35 Gy/hr (Fig. 3A,C). In response to stress such as IR, *A. pullulans* turns black due to the production of melanin (Gostinčar et al., 2014). Exposing a partially lead-shielded *A. pullulans* plated culture to IR caused the exposed half to produce melanin and turn black (Fig. 3D, E). IR-exposed hyphae accumulated melanin in the hyphal tips (Fig. 3F vs G, arrow). The black, melanized culture regenerated over time, with newly formed hyphae from fully exposed or shielded colonies growing white, indicating the non-melanized form replenished growth (Fig. 3H,I, arrowheads).

### *Drosophila* tolerate a diet of highly radioresistant fungi

The preferred diet of *Drosophila* species is yeast growing on various organic matter, including fruit (Hoang et al., 2015; Meshrif et al., 2016). In a lab setting, *Drosophila melanogaster* prefers *Saccharomyces cerevisiae* (*S. cerevisiae*). To determine whether radiation-resistant fungi are a preventative dietary approach against IR, we first tested whether *Drosophila* tolerated a diet of exclusive *R. taiwanensis* or *A. pullulans* (Fig. 4). To feed flies both fungal strains, we harvested the fungi and placed a smear on a small agar plate. Flies fed *ad libitum* on the fungi while placed in a standard egg-laying cup with the plate on the bottom (Fig. 4A). Both larvae and adult flies ingested *R. taiwanensis* as indicated by their red guts (Fig. 4B,C, arrows). To better visualize ingested *A. pullulans*, we fed the flies irradiated, melanized fungi. The black fungi could also be seen in the larval and adult fly guts (Fig. 4D,E, arrows). This indicated that the flies readily consumed both fungal strains. We also examined the sensitivity of flies at developmental stages to determine whether fungal feeding affects pupation and eclosion rates (Fig. 4F). All larvae developed normally into pupae and eclosed at the same rate compared to control (*S. cerevisiae*) regardless of the kind of fungus (Fig. 4F). In addition, the flies survived for weeks exclusively eating either fungus (Fig. 4G). Flies fed on both strains had a reduced lifespan compared to those fed exclusive *S. cerevisiae* (Fig. 4G, blue and green vs black lines), with *A. pullulans* having a greater effect (Fig. 4G). However, neither was acutely toxic, with flies able to survive approximately 30 or 50 days, respectively, for *A. pullulans* or *R. taiwanensis*.

**Fig. 4:**
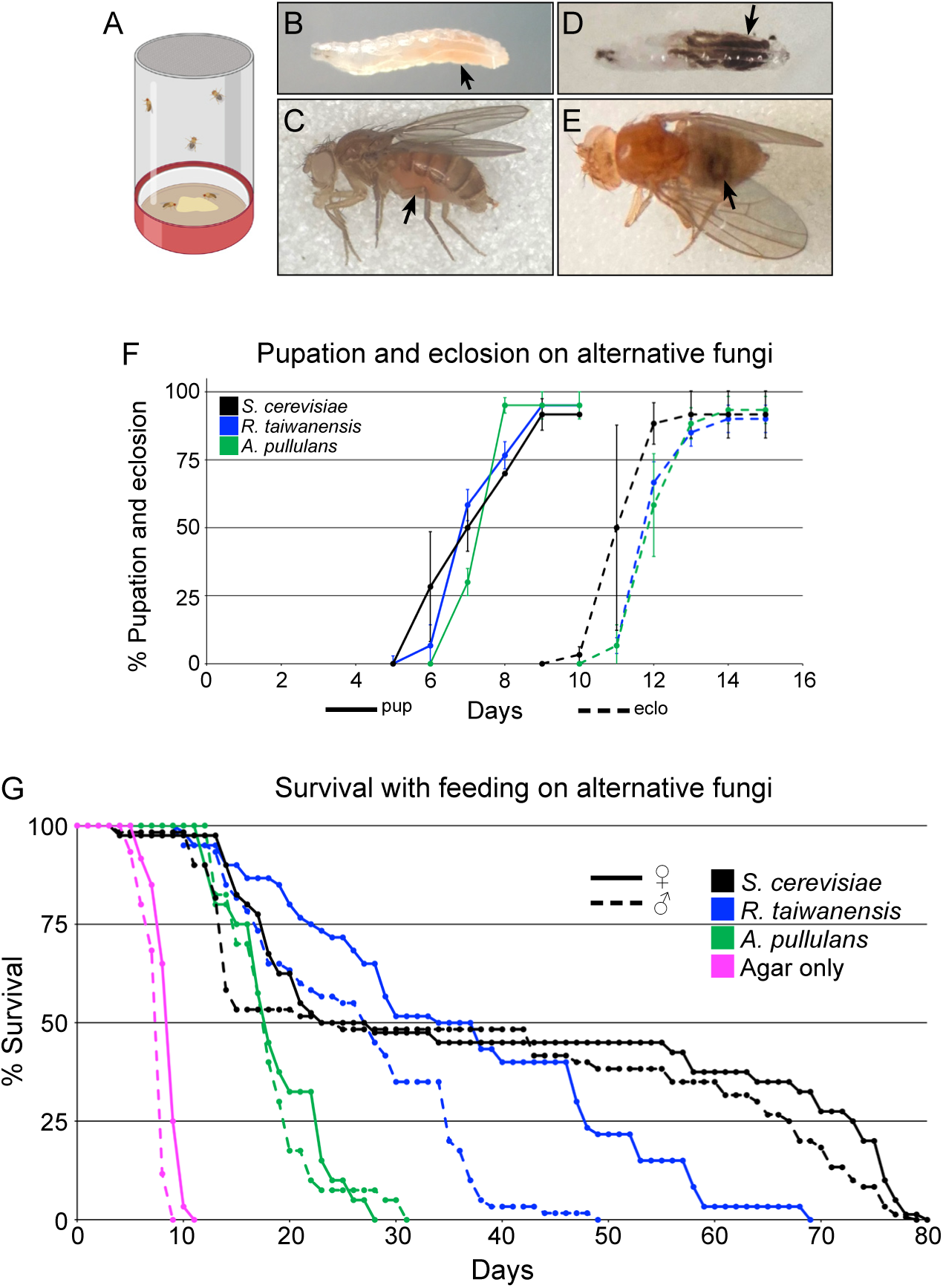
*Drosophila* tolerate dietary *Aureobasidium pullulans* and *Rhodotorula taiwanensis*. (A) Schematic of egg laying cups. Harvested *R. taiwanensis* and *A. pullulans* were placed on an agar plate for fly consumption (bottom). (B, C) Larva (B) and adult female fly (C) after feeding on *R. taiwanensis.* Red guts (arrows) indicate the animals ingested the fungi. (D, E) Larva (D) and adult female fly (E) after feeding on irradiated *A. pullulans.* Guts show the animals ingested the radiation-induced melanized fungi (arrows). (F) The percentage of larvae that pupated (solid lines) and eclosed (dashed lines) after being fed exclusively on alternative fungi. Each point on the graph represents the average of triplicates. Error bars represent standard deviation. (G) Lifespan analysis of males and females exclusively fed *A. pullulans* or *R. taiwanensis*. Agar only plates had no corn syrup (pink lines). Graphs were plotted using Microsoft Excel. Each point on the lifespan represents the average of triplicates of twenty flies that were fed fungi in parallel.

### Feeding *A. pullulans*, but not *R. taiwanensis*, improves male survivorship after acute irradiation

Previous research suggested that radiation-induced metabolic changes or antioxidant production may endow fungi with radioresistance (Dadachova et al., 2007; Jung et al., 2016; Kelley et al., 2022). To test whether pre-feeding *Drosophila* radioresistant fungus could protect fly lifespan after acute IR exposure, we fed newly eclosed males or female for two days before exposure to acute IR, then monitored the flies for their entire lifespan (Fig. 5A). We found that flies fed *A. pullulans* showed a sex-specific improvement to lifespan, with males having increased lifespan, but females not (Fig. 5B, C). In contrast, neither males nor females had increased lifespan with *R. taiwanensis* pre-feeding (Fig. 5D,E). Consistent with the exclusive diet of *R. taiwanensis* (Fig. 4G), females exhibited decreased lifespan (Fig. 5E, green and orange lines). Since fungal exposure to IR upregulates protective cellular components, we next tested whether IR-exposed, melanized fungi conferred greater dietary protection against acute IR. For *A. pullulans*, we found that males still exhibited lifespan improvement (Fig. 5B, p = 0.0166 (+ γ, magenta) vs p = 0.0148 (- γ, blue)), but this effect was not enhanced with the melanized fungus. Females again did not show any improvement in lifespan by feeding melanized *A. pullulans* (Fig. 5C). Feeding irradiated *R. taiwanensis* did not increase lifespan in either males or females and appeared to instead make the fungus more toxic (Fig. 5D (+ γ, magenta),E (+ γ, orange)).

**Fig. 5:**
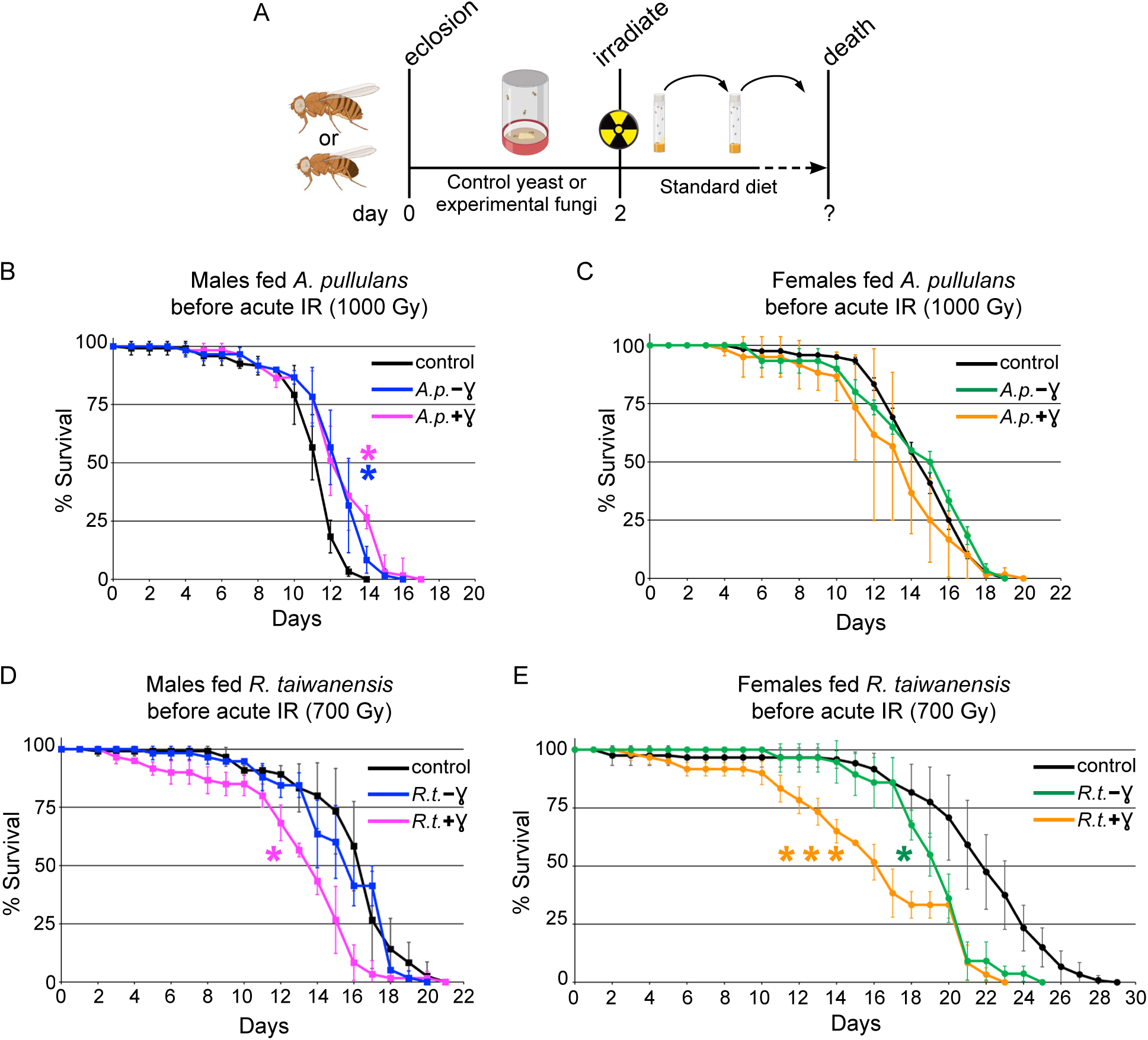
Feeding *Aureobasidium pullulans* improves lifespan in males. (A) Schematic showing the experimental timeline. Newly eclosed males and females are fed control yeast or experimental fungal paste for two days. The flies were subsequently placed in standard food vials and irradiated. Flies were transferred to fresh vials every 1-2 days until all died. (B) Lifespan of males fed non-irradiated *A. pullulans* (*A.p.*) (blue, −ɣ) and chronically irradiated *A. pullulans* (magenta, +ɣ) before exposure to 1000 Gy acute irradiation. (C) Lifespan of females fed non-irradiated *A. pullulans* (green, −ɣ) and chronically irradiated *A. pullulans* (orange, +ɣ) before exposure to 1000 Gy acute irradiation. Prophylactic dietary *A. pullulans* significantly extended the lifespan of males, but not females. (D) Lifespan of males fed non-irradiated *R. taiwanensis* (*R.t.*) (blue, −ɣ) and chronically irradiated *R. taiwanensis* (magenta, +ɣ) before exposure to 700 Gy acute irradiation. Feeding irradiated *R. taiwanensis* was detrimental to male lifespan. (E) Lifespan of females fed non-irradiated *R.t*. (green, −ɣ) and chronically irradiated *R. taiwanensis* (orange, +ɣ) before exposure to 700 Gy acute irradiation. Feeding non-irradiated and irradiated *R. taiwanensis* to females significantly decreased their lifespan. Graphs were plotted using Microsoft Excel. Each point on the lifespan represents the average of triplicates of twenty flies that were irradiated simultaneously. Error bars represent standard deviation. Statistical analysis was calculated using Online Application for Survival Analysis 2 (OASIS) and statistical significance was calculated using the Wilcoxon-Breslow-Gehan test. P values vs control are as follows: (B) p = 0.0331 (*) for *A.p.*(+ɣ), 0.0297 (*) for *A.p.*( −ɣ) ; (C) p=0.461 for *A.p.*(+ɣ), 1.0 for *A.p.*(−ɣ) (D) p= 0.0041 (*) for *R.t.*(+ɣ), 1.0 for *R.t.*(−ɣ) (E) p= 0.0002 (***) for *R.t.*(+ɣ), 0.0116 (*) for *R.t.*(−ɣ).

### Feeding *A. pullulans* attenuates IR-induced cellular damage in male guts

Since male guts were more susceptible to IR exposure (Fig. 2) and pre-feeding with *A. pullulans* improved male lifespan (Fig. 5B), we examined whether *A. pullulans* pre-feeding could ameliorate any negative effects of IR on gut morphology (Fig. 6). Feeding male fruit flies *A. pullulans* appeared to have a dual effect on their gut enterocytes. While it reduced the number of enterocytes with abnormal nuclei (Fig. 6C,C”,E, Fig. S2G,G”,H,H”,I,I”), *A. pullulans* feeding without radiation disrupted the cells’ actin cytoskeleton, leading to compromised cellular barriers and altered cell shape (Fig. 6C,C’,F, Fig. S2G,G’H,H’,I,I’). This disruption might explain the observed shortened lifespan (Fig. 4G). However, when the *A. pullulans*-fed flies were exposed to radiation, they exhibited fewer abnormal nuclei in their enterocytes compared to flies that were irradiated without dietary *A. pullulans* (Fig. 6B, B”vs D,D”,E). Even though radiation still increased the disruption of cell barriers in the *A. pullulans*- fed group (Fig. 6D,D’,F, SFig. 2J,J’,K,K’,L,L’), these findings collectively support that feeding male *Drosophila* the radioresistant *A. pullulans* helped lessen or postpone gut injury, ultimately improving their survival following radiation exposure.

**Fig. 6:**
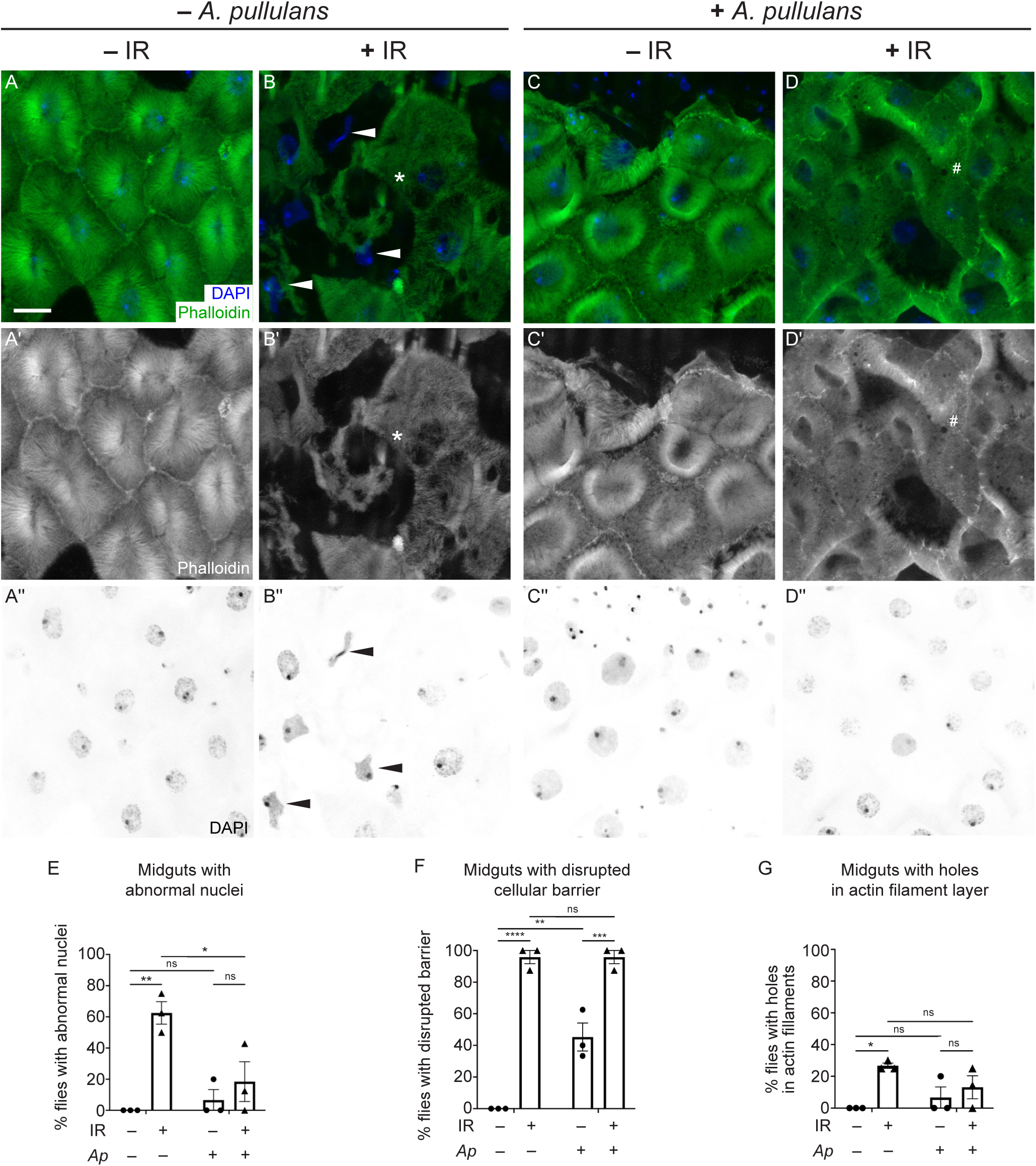
Feeding *Aureobasidium pullulans* attenuates IR-induced cellular damage in male guts. (A-D”) Dissected midguts from day 4 adult males exposed to IR at day 2 and control males labeled with phalloidin to label actin filaments and DAPI to label nuclei. The images of the R4 regions of midguts from each experimental condition were obtained using a confocal microscope with z-stacked for a layer of enterocytes. (A-A”) Representative gut dissected from a male control, showing clear cell barriers (A,A’) and normal nuclear shapes in enterocytes (A,A”). (B-B”) Representative gut dissected from a male two days after irradiation. Enterocytes had aberrant nuclear shape (B, B”, arrowheads, E) and altered cellular morphology (B,B’,F). Holes were formed within a layer of actin filament (B,B’, asterisk, G) by irradiation. (C-C”) Representative gut from a male fed *A. pullulans* (AP) had normal nuclear morphology (C,C”,E) in enterocytes while having a loss of cell barriers with some degree (C,C’,F). (D-D”) Representative gut from a male fed *A. pullulans* followed by irradiation had normal nuclear shape (D,D”,E) as well as loss of cellular barriers and irregular cell shape (D,D’,F). Many small holes appeared within actin layers (D,D’, number sign) but were not counted because they did not satisfy the criterion as described in Methods. Each graph indicates the percentage of animals having guts with abnormal nuclei (E), (F) disrupted cellular barriers (F), or holes in the actin filaments layer in enterocytes (G). Quantification method is described in the Materials and Methods section. Each point on the graph represents the percentage of each replicate, and each bar presents the arithmetic mean of triplicates with a standard error of the mean (SEM). Data were analyzed statistically using a two-way ANOVA with multiple comparisons, followed by Tukey’s post hoc analysis. The following numbers of the midguts were assayed per experiment: male control (11, 6, 7); male +IR (8, 4, 10); male fed *A.p.* (8, 6, 5); male fed *A.p.*-IR (7, 8, 7). *P* values of multiple comparisons are as follows: (E) p = 0.0026 (**) for (-/+IR) without *A.p.* and 0.0197 (*) for (- /+*A.p.*) with IR; (F) p < 0.0001 (****) for (-/+IR) without *A.p.*, 0.0007 (***) for (-/+IR) with *A.p.*, and 0.0014 (**) for (-/+*A.p.*) without IR; (G) p = 0.0225 (*) for (-/+IR) without *A.p*. ns = not significant. (A,B,C,D) Green = phalloidin, blue = DAPI. (A’,B’,C’,D’) White = phalloidin. (A”,B”,C”,D”) White = DAPI. (A-G) -IR: non-irradiated, +IR: 1000 Gy irradiation. (E,F,G) – *A.p.*: non-feeding *A. pullulans*, +*Ap*: feeding *A. pullulans*. Scale bar: 10 μm in A for A-D”.

## Discussion

### Identifying palliative strategies for radiation exposure

Radiotherapy patients, particularly those exposed to abdominal and pelvic radiation therapy, frequently have the side effect of GI tract damage and enteropathy (Fernandes & Andreyev, 2021; Hauer-Jensen et al., 2014). An important consideration is that symptoms may be delayed and impact long-term quality of life. In addition to tissue damage, one notable change in this cohort is the alteration in the microbiome (Oh et al., 2021). Changes to metabolites, tryptophan pathway intermediates in particular, and the microbiome occur in mice that exhibit improved survivorship to irradiation as well as in patients with leukemia who experienced fewer adverse effects of whole-body irradiation (Guo et al., 2020). Identifying protective measures would greatly aid these patients.

### *Drosophila* as a model to study dietary protection from IR

Drosophila has many advantages for evaluating preventative measures against radiation damage compared to mammalian models, such as experimental numbers in the hundreds, longitudinal analyses to capture survival, developmental, reproductive, and aging metrics, as well as observations of delayed effects of both radiation injury and treatment. Since irradiators can vary sample exposures due to differences in source and geometry, we created the survival curves for male and female lifespan using our ^137^Cs source, ensuring the flies were at a consistent and correct position in the irradiator and that six vials of adults, three experimental and three control, could be simultaneously exposed. As demonstrated in many species, we found sex-specific IR sensitivity in Drosophila, with males being more sensitive than females and thus having shorter lifespans (Gartner, 1973). This sex-specific difference occurred at all doses we tested. In general, female flies are more stress-tolerant than males (Pomatto et al., 2018).

The survival radiosensitivity we observed with males compared to females was reflected in their gut morphologies, with males experiencing more damage. The GI tract is highly conserved in function and structure between Drosophila and vertebrates (Capo et al., 2019; Douglas, 2018; Ludington & Ja, 2020). The fly intestinal epithelium has become a model to study tissue repair and age-related decline of tissue regeneration, including response to IR exposure (Biteau et al., 2008; Guo et al., 2016; Sharma et al., 2020). Intestinal stem cells (ISCs) are the primary cells that divide to respond to damage of the intestinal epithelium. Normally quiescent, ISCs proliferate in response to oxidative stress (Biteau et al., 2008). Epithelial dysplasia and barrier dysfunction develop in flies subjected to oxidative stresses such as aging, radiation, and ROS- generating compounds, and genetic disruptions of mitochondrial function induce epithelial barrier dysfunction (Rera et al., 2011; Rera et al., 2012; Sharma et al., 2020). Since pre-feeding *A. pullulans* improved male gut morphology after IR exposure and given the high degree of conservation in GI architecture and signaling pathways between Drosophila and mammals, dietary *A. pullulans* represents a potential nutritional preventative measure that may directly act to protect the GI tract from IR-induced injury.

### Mechanisms and molecules for IR protection

IR causes injury directly and indirectly. Direct cell and tissue injury occurs from IR damage to DNA, proteins and lipids, including DNA repair enzymes. The indirect damage is due to water radiolysis that increases the concentration of free radicals which subsequently damage cellular components. Metazoan strategies of differential tolerance to the effect of IR may be due to both better DNA protection and increased ability to handle reactive oxygen species (Kelleher et al., 2025). How might *A. pullulans* exert its protective effect on the gut? One possibility is that ingested antioxidants could be taken up by the enterocytes, thus increasing protection against free radicals. For example, feeding mice melanin-containing Jelly Ear mushrooms compared to porcini mushrooms greatly improved survival post 9 Gy IR, with no signs of GI radiation damage 45 days post-irradiation (Revskaya et al., 2012). Survival improvement post-irradiation has also been demonstrated with the oral administration of antioxidants (Brown et al., 2010; Ghosh et al., 2009). N-acetyl-L-cysteine as a dietary supplement protected the mouse intestinal epithelial barrier after irradiation (Shukla et al., 2016). Mice ingesting *Aloe vera* extract had delayed radiation sickness symptoms and lower acid phosphatase and alkaline phosphatase levels in the liver after 6 Gy irradiation compared to a control (Gehlot et al., 2010). A mixture of ingested traditional Indian medicinal plants increased survival and improved DNA damage compared to a control after 7.5 Gy exposure (Sandhya et al., 2006). In Drosophila, feeding curcumin, a plant phenolic compound, increased lifespan and decreased protein carbonylation (Seong et al., 2015). Feeding flies a tea polyphenol and beta-carotene decreased mutation frequency and increased antioxidant levels post 10 Gy exposure compared to controls (Nagpal & Abraham, 2017). Finally, flies fed ibuprofen or two flavonoids (quercetin and epicatechin) showed increased lifespan compared to a control after 1000 Gy exposure (Proshkina et al., 2016).

Pre-feeding MnCl_2_ before IR exposure increased radiation survival in a similar manner to *A. pullulans* feeding (Volpe et al., 2025). The protective effect of dietary manganese was attributed to an increase in small molecule manganous peptide antioxidant content, not an increase in the free radical scavenger Mn Superoxide Dismutase. It is possible that *A. pullulans* contains high levels of manganous peptides, and this could explain its protective effect. Melanin is recognized as a radioprotectant present in extremophile microorganisms and has been shown to have radioprotective effects in vertebrates as well (Kunwar et al., 2012; Le Na et al., 2019; Revskaya et al., 2012). Mice fed melanin-containing mushrooms had greater gastrointestinal protection compared to mice fed non-melanized mushrooms, supporting that the presence of melanin in the gut could protect the gut lining from the effects of acute irradiation (Revskaya, 2012). *A. pullulans* melanizes in response to IR (Campana et al., 2022).

Since this fungus extended lifespan in irradiated males, we also tested the melanized phenotype to determine whether melanin could increase its effectiveness. However, melanized *A. pullulans* did not increase the protective effect suggesting that the concentration of melanin was not sufficient to produce an enhanced radioprotective effect or that other metabolites or molecules are responsible for the protection.

### Wider implications of identifying dietary strategies for reducing radiation-induced cellular damage

Radiotherapy patients are the most recognizable cohort that would benefit from identifying new strategies for reducing the effects of radiation injury. An additional population at serious health risk from IR are those who are potentially exposed to radiation disasters – either as cleanup workers, military personel, or the general public. For these groups of people, the diverse and wide-ranging effects of IR exposures are organized into subsyndromes of acute radiation syndrome (ARS). While medical countermeasures to the hematopoietic subsyndrome of acute radiation syndrome (H-ARS) have been developed, no medical countermeasures to the gastrointestinal subsyndrome (GI-ARS) exist. There are only four countermeasures approved as mitigators by the FDA for H-ARS, but none are effective as prophylactic intervention or for GI damage (Bene et al., 2021). Gamma tocotrienol, a naturally occurring vitamin E isoform, reportedly confers some radioprotective survival in mice (Ghosh et al., 2009; Kumar et al., 2020; Singh & Seed, 2020). Effective prophylactic countermeasures targeting the vulnerable gastrointestinal tract need to be identified (Singh & Seed, 2017, 2020).

### Future directions and unanswered questions

Here, we identified *A. pullulans* as a dietary prevention strategy for protection against acute irradiation. This observation opens several future directions. If future vertebrate studies indicate that dietary *A. pullulans* can protect against radiation injury, it could prove beneficial to humans. We have only observed that feeding the entire fungus conveys a protection, however, in the future the exact compound(s) could be identified. Finally, there are many radioresistant fungi that could be tested in this manner. While some contributors to their radioresistance have been identified, such as manganous peptides, there are likely other unidentified mechanisms and metabolites at work. Using whole fungi that have adapted to survival after high doses of irradiation may be the best way to protect people from the deleterious effects of IR.

## Acknowledgments

This work was supported by the National Institutes of Health R21OD034471 and R01GM127938 to R. T. C.

## Author Contributions

Conceptualization, R.P.V. and R.T.C.; methodology, R.P.V., H.J.H., and R.T.C.; validation, R.P.V. and H.J.H.; formal analysis, R.P.V and H.J.H.; writing—original draft preparation, R.P.V. and R.T.C..; writing—review and editing, R.P.V., H.J.H. and R.T.C.; supervision, R.T.C.; funding acquisition, R.T.C. All authors have read and agreed to the published version of the manuscript.

## Conflicts of interest

The authors declare no competing interests.

## Disclaimer

The funders had no role in the design of the study; in the collection, analyses, or interpretation of data; in the writing of the manuscript, or in the decision to publish the results. The content is solely the responsibility of the authors and does not necessarily represent the official views of Uniformed Services University, the Department of Defense, the National Institutes of Health or the Henry M. Jackson Foundation for the Advancement of Military Medicine, LLC.

## Notes

### Competing Interest Statement

The authors have declared no competing interest.

