## Supplementary material for "Feeding Drosophila highly radioresistant fungi improves survival and gut morphology following acute gamma radiation exposure": Figure S1

Figure S1. Drosophila male guts are more sensitive to the effects of irradiation compared to female

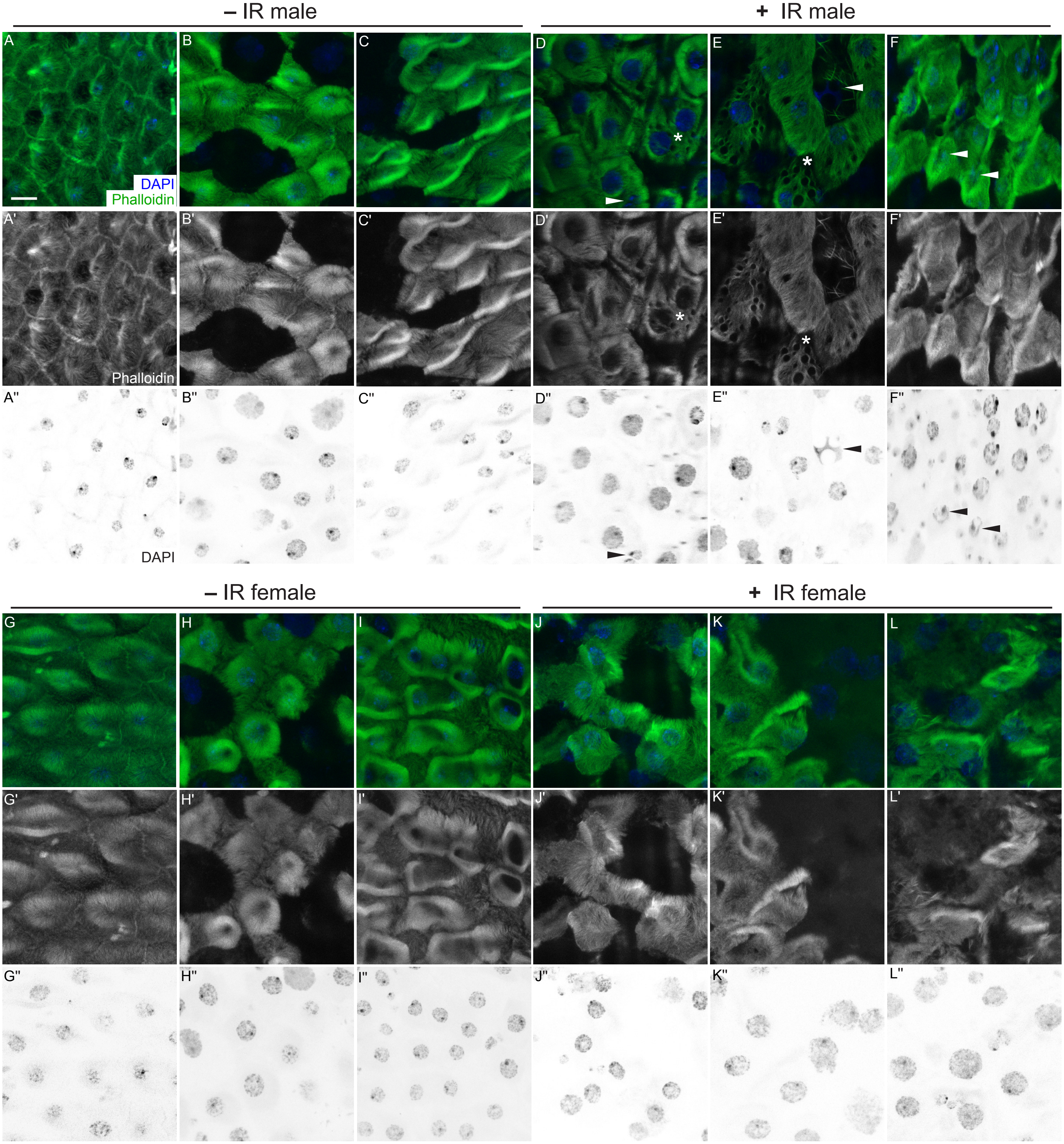

**Supplementary Figure 1. *Drosophila* male guts are more sensitive to the effects of irradiation compared to females.**

(A-C”) R4 regions of midguts from male controls and (G-I”) female controls. Dissected midguts were labeled with phalloidin to label actin filaments and DAPI to label nuclei, and then imaged using a confocal microscope with z-stacks for a layer of enterocytes. The enterocytes had clear cellular barriers in both males (A,A’,B,B’,C,C’) and females (G,G’,H,H’,I,I’) as well as normal nuclear shape (A,A”,B,B”,C,C”, male; G,G”,H,H”,I,I”, female). (D-F”) R4 regions of midguts from males two days after irradiation. Immunostaining shows disrupted cellular barriers (D,D’E,E’F,F’), multiple holes in actin filament layers (D,D’E,E’, asterisks), or abnormal nuclear shape (E,E”,F,F”, arrowheads). (J-L”) R4 regions of midguts from females two days after irradiation. Immunostaining shows disrupted cellular barriers (J,J’,K,K’,L,L’), but relatively normal nuclear shape (J,J”,K,K”,L,L”). (A-L”) -IR: non-irradiation, +IR: 1000 Gy irradiation. (A,B,C,D,E,F,G,H,I,J,K,L) Green = Phalloidin, blue = DAPI. (A’,B’,C’,D’,E’,F’,G’,H’,I’,J’,K’,L’) White = phalloidin. (A”,B”,C”,D”,E”,F”,G”,H”,I”,J”,K”,L”) White = DAPI. Scale bar: 10 µm in A for A-L”.
