## Supplementary material for "Feeding Drosophila highly radioresistant fungi improves survival and gut morphology following acute gamma radiation exposure": Figure S2

### Figure S2. Prophylactic dietary *A. pullulans* mitigates IR-induced cellular damage in male guts

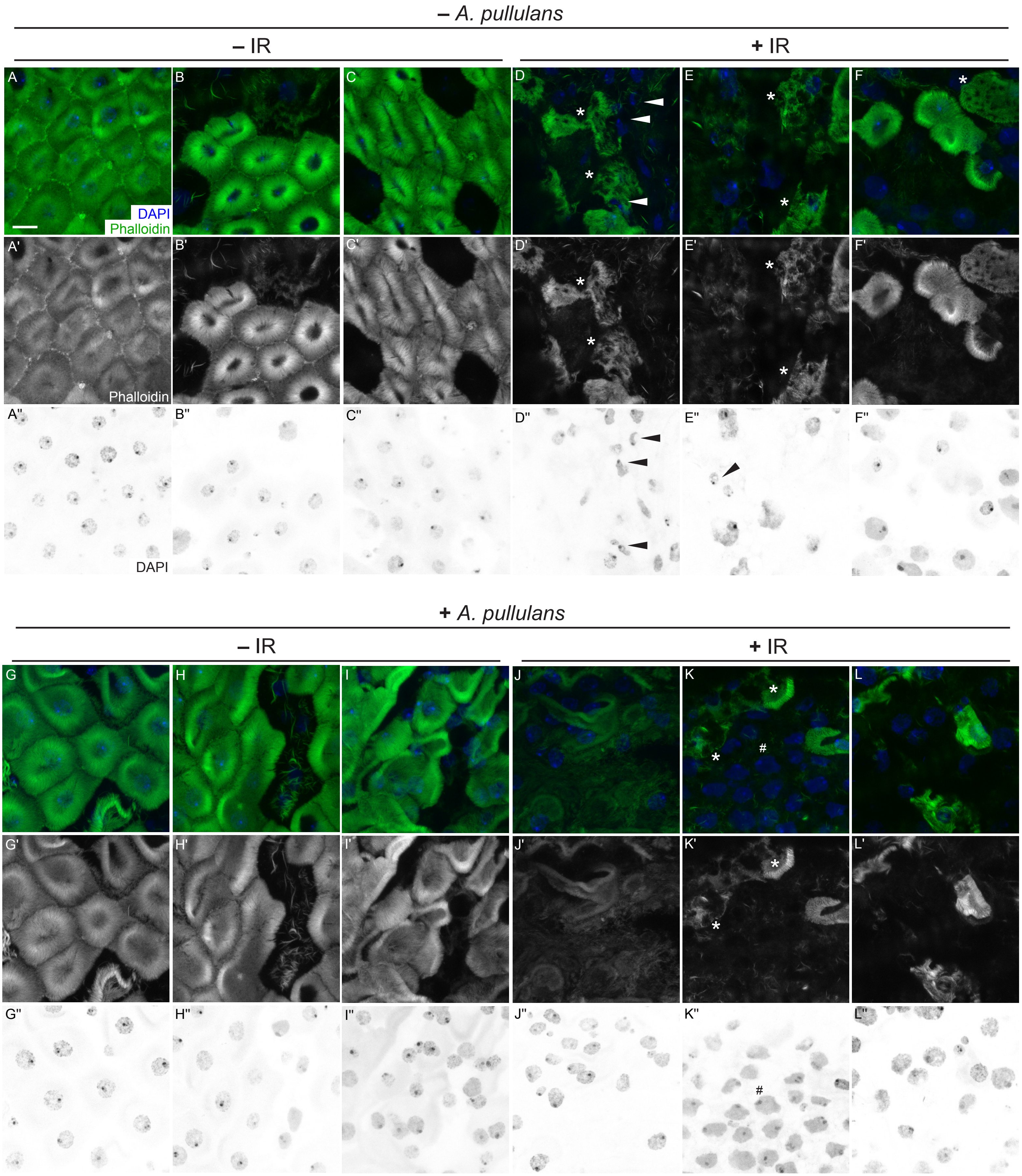

**Supplemental Figure 2. Prophylactic dietary *Aureobasidium pullulans* mitigates IR-induced damaging process of male guts.**

(A-C”) R4 regions of midguts from male controls. Dissected midguts were labeled with phalloidin to label actin filaments and DAPI to label nuclei, and then imaged using a confocal microscope with z-stacks for a layer of enterocytes. The enterocytes had clear cellular barriers (A,A’,B,B’,C,C’) and normal nuclear shape (A,A”,B,B”,C,C”). (D-F”) R4 regions of midguts from males two days after irradiation. Immunostaining shows abnormal nuclear shape (D,D”,E,E”), disrupted cellular barriers (D,D’E,E’F,F’), and holes within an actin filament layer (D,D’E,E’F,F’, asterisk) (G-I”) R4 regions of midguts from males fed *Aureobasidium pullulans* (*A. pullulans*)**.** Feeding *A. pullulans* caused some degrees of loss of cellular barriers (G,G’,H,H’,I,I’). (J-L”) IR following *A. pullulans* feeding did not induce smaller nuclei although nuclear morphology was altered (K,K”, number sign). Actin filament structure was severely changed by IR (J,J’,K,K’,L,L’). (A-L”) -IR: non-irradiation, +IR: 1000 Gy irradiation, - *A. pullulans*: non-fungus feeding, + *A. pullulans*: fungus feeding. (A,B,C,D,E,F,G,H,I,J,K,L) Green = Phalloidin, blue = DAPI. (A’,B’,C’,D’,E’,F’,G’,H’,I’,J’,K’,L’) White = phalloidin. (A”,B”,C”,D”,E”,F”,G”,H”,I”,J”,K”,L”) White = DAPI. Scale bar: 10 µm in A for A-L”.
