## Supplementary material for "Feeding Drosophila highly radioresistant fungi improves survival and gut morphology following acute gamma radiation exposure": Table S1

**Supplementary Table 1.** Influence of Dietary Fungi on Radiation Survival

| Group | Dose | Diet | Mean | S.E. | Log Rank Bonferroni P-value | | | Gehan-Breslow-Wilcoxon P-value | | |
|---|---|---|---|---|---|---|---|---|---|---|
| | | | | | P (vs Control) | P (vs Aureo+Mel) | P (vs Aureo-Mel) | P (vs Control) | P (vs Aureo+Mel) | P (vs Aureo-Mel) |
| Female | 1000 Gy | Control | 15.85 | 0.5 | - | 1 | 1 | - | 0.461 | 1 |
| Female | 1000 Gy | Aureobasidium(+Mel | 14.65 | 0.74 | 1 | - | 0.8718 | 0.461 | - | 0.6429 |
| Female | 1000 Gy | Aureobasidium(-Mel) | 15.55 | 0.73 | 1 | 0.8719 | - | 1 | 0.6429 | - |
| Male | 1000 Gy | Control | 12.45 | 0.34 | - | 0.0086 | 0.0142 | - | 0.0331 | 0.0297 |
| Male | 1000 Gy | Aureobasidium(+Mel | 13.9 | 0.48 | 0.0086 | - | 0.8374 | 0.0331 | - | 1 |
| Male | 1000 Gy | Aureobasidium(-Mel) | 13.65 | 0.41 | 0.0142 | 0.8374 | - | 0.0297 | 1 | - |
| **Group** | **Dose** | **Diet** | **Mean** | **S.E.** | **Log Rank Bonferroni P-value** | | | **Gehan-Breslow-Wilcoxon P-value** | | |
| | | | | | **P (vs Control)** | **P (vs Rhodo+ɣ)** | **P (vs Rhodo-ɣ)** | **P (vs Control)** | **P (vs Rhodo+ɣ)** | **P (vs Rhodo-ɣ)** |
| Female | 700 Gy | Control | 23.3 | 0.75 | - | 0.0001 | 0.0062 | - | 0.0002 | 0.0116 |
| Female | 700 Gy | Rhodotorula(+ɣ) | 17.45 | 1.04 | 0.0001 | - | 0.2444 | 0.0002 | - | 0.0783 |
| Female | 700 Gy | Rhodotorula(-ɣ) | 20.6 | 0.54 | 0.0062 | 0.2444 | - | 0.0116 | 0.0783 | - |
| Male | 700 Gy | Control | 17.4 | 0.57 | - | 0.0143 | 1 | - | 0.0041 | 1 |
| Male | 700 Gy | Rhodotorula(+ɣ) | 14.45 | 0.84 | 0.0143 | - | 0.0664 | 0.0041 | - | 0.0407 |
| Male | 700 Gy | Rhodotorula(-ɣ) | 16.85 | 0.63 | 1 | 0.0664 | - | 1 | 0.0407 | - |

* for average of n=20 triplicate
